# PQS-Induced Outer Membrane Vesicles Enhance Biofilm Dispersion in *Pseudomonas aeruginosa*

**DOI:** 10.1101/2020.06.15.153908

**Authors:** Adam C. Cooke, Catalina Florez, Elise B. Dunshee, Avery D. Lieber, Michelle L. Terry, Caitlin J. Light, Jeffrey W. Schertzer

**Author notes:** Address correspondence to Jeffrey W. Schertzer. These authors contributed equally to this work (Adam C. Cooke, Catalina Florez). Author order was determined alphabetically.

## Abstract

Bacterial biofilms are major contributors to chronic infections in humans. Because they are recalcitrant to conventional therapy, they present a particularly difficult treatment challenge. Identifying factors involved in biofilm development can help uncover novel targets and guide the development of anti-biofilm strategies. *Pseudomonas aeruginosa* causes surgical site, burn wound, and hospital acquired infections, and is also associated with aggressive biofilm formation in the lungs of cystic fibrosis patients. A potent but poorly understood contributor to *P. aeruginosa* virulence is the ability to produce outer membrane vesicles (OMVs). OMV trafficking has been associated with cell-to-cell communication, virulence factor delivery, and the transfer of antibiotic resistance genes. Because OMVs have almost exclusively been studied using planktonic cultures, little is known about their biogenesis and function in biofilms. Our group has shown that the Pseudomonas Quinolone Signal (PQS) induces OMV formation in *P. aeruginosa*, and in other species, through a biophysical mechanism that is also active in biofilms. Here, we demonstrate that PQS-induced OMV production is highly dynamic during biofilm development. Interestingly, PQS and OMV synthesis are significantly elevated during dispersion, compared to attachment and maturation stages. PQS biosynthetic and receptor mutant biofilms were significantly impaired in their ability to disperse, but this phenotype could be rescued by genetic complementation or exogenous addition of PQS. Finally, we show that purified OMVs can actively degrade extracellular protein, lipid, and DNA. We therefore propose that enhanced production of PQS-induced OMVs during biofilm dispersion facilitates cell escape by coordinating the controlled degradation of biofilm matrix components.

**Importance:** Treatments that manipulate biofilm dispersion hold the potential to convert chronic drug-tolerant biofilm infections from protected sessile communities into released populations that are orders-of-magnitude more susceptible to antimicrobial treatment. However, dispersed cells often exhibit increased acute virulence and dissemination phenotypes. A thorough understanding of the dispersion process is therefore critical before this promising strategy can be effectively employed. PQS has been implicated in early biofilm development, but we hypothesized that its function as an OMV inducer may contribute at multiple stages. Here, we demonstrate that PQS and OMVs are differentially produced during *Pseudomonas aeruginosa* biofilm development and that effective biofilm dispersion is dependent on production of PQS-induced OMVs, which likely act as delivery vehicles for matrix degrading enzymes. These findings lay the groundwork for understanding the roles of OMVs in biofilm development and suggest a model to explain the controlled matrix degradation that accompanies biofilm dispersion in many species.

## Introduction

It has long been appreciated that biofilms contribute to a majority of bacterial infections (1–4). Biofilm cells differ from planktonic cells in phenotype (5), gene expression (6), and protein production (7–10). These differences provide biofilm cells enhanced tolerance to antibiotics and host defenses (11–14). *Pseudomonas aeruginosa* is a clinically relevant and highly studied model organism for biofilm development. Surface-attached *P. aeruginosa* biofilms develop in a stepwise fashion where bacteria first reversibly and then irreversibly attach to a surface (7). The maturation phase is marked by the emergence of three-dimensional microcolonies during maturation I and the formation of mushroom-like clusters during maturation II (7). In response to external or endogenous cues, the final phase is initiated when bacterial cells erupt from the biofilm and disperse (7). During dispersion, motile bacteria degrade the extracellular polymeric matrix that encases them, colonize new surfaces, and recommence the biofilm life cycle (7, 15). Identification of the factors that regulate biofilm development is essential for the creation of novel therapeutics against these recalcitrant bacterial communities.

Quorum signaling is known to regulate *P. aeruginosa* biofilm formation (7, 16). Specifically, the Las system controls the progression from reversible to irreversible attachment (16), and the Rhl system controls the transition from irreversible attachment to maturation I (7). The Pseudomonas Quinolone Signal (PQS) has also been proposed to regulate biofilm development (17, 18). Production of PQS is initiated by the Las system through direct activation of the genes encoding the PQS regulator, PqsR (18, 19), and the biosynthetic FAD-dependent monooxygenase, PqsH (20, 21). PQS controls the production of many virulence factors (17), including elastase, pyocyanin (22), and iron chelators (23–25). It has been reported that PQS biosynthetic mutants are deficient in the formation of mushroom-shaped microcolonies, which are characteristic of mature biofilms (26, 27). Several hypotheses aim to connect the contributions of PQS in biofilm development to its functionality as a cell-to-cell communication signal. Rampioni and coworkers (28) suggested that PQS controls biofilm development via PqsE-dependent signaling, activating the Rhl system and its downstream effectors. It has also been shown that extracellular DNA (eDNA) contributes to biofilm maturation and that PQS-induced prophage activation results in DNA release into the biofilm (26). The buildup of HQNO, which is controlled by PQS signaling, likewise results in autolysis, eDNA release, and increased biofilm biomass (29). We were interested in exploring whether other well-documented functions of PQS may also play a role during the various stages of biofilm development.

In addition to its role as a signaling molecule, PQS is also known to modulate production of outer membrane vesicles (OMVs) (30–34). OMVs are spherical structures derived from the outer membrane of Gram-negative bacteria that range from 50-300 nm in diameter (35–38). These nanostructures form a dedicated transport system that helps deliver cell-to-cell communication signals (30, 39, 40), nucleic acids (41, 42), proteases (43, 44), antibiotic degrading enzymes (45, 46), lytic enzymes (47–49), iron chelators (23–25), and antibiotic resistance genes (50). In conjunction with their function as transport machinery, OMVs have also been associated with biofilm development in *Helicobacter pylori* (51), *Vibrio cholerae* (52), and *Pseudomonas putida* (53). Little is known about the roles that OMVs play in *P. aeruginosa* biofilms. However, it has been reported that OMVs are commonly found within biofilms produced by this organism (44, 54) and that their production is controlled by PQS (55).

PQS induces OMV production through a biophysical mechanism that is driven by favorable interactions with lipopolysaccharide (LPS) in the outer leaflet of the outer membrane (OM) (32, 56). These interactions promote asymmetric expansion of the outer membrane, which induces membrane curvature and ultimately leads to the production of OMVs (33). The importance of PQS in OMV production is evident from many experiments involving deletions in early biosynthetic genes (e.g. *pqsA*, coding for the anthrailoyl-CoA ligase responsible for the first step in alkyl-quinolone biosynthesis (57–59)), late biosynthetic genes (e.g. *pqsH*, coding for the flavin-dependent monooxygenase responsible for the final step in PQS biosynthesis (20, 21, 60, 61)), and the PQS receptor (*pqsR*) (19, 62). Deletion of any of these genes results in drastic reductions or outright abrogation of OMV biogenesis in planktonic cultures. Our recent work demonstrated that loss of PQS production also compromised OMV production in *P. aeruginosa* biofilms (55). Importantly, use of these well-characterized mutants (in addition to others such as *pqsE*) can help detangle the biophysical roles of PQS from its role as a signaling molecule, as well as clarify contributions directly related to PQS from those of other related alkyl-quinolones. While several studies have implicated PQS in the development of *P. aeruginosa* biofilms, it is not known if PQS is involved all stages of biofilm formation. Additionally, it remains unclear if PQS affects biofilm development due to its role in quinolone signaling, virulence factor production, OMV biogenesis, or any combination of these. The current study presents a comprehensive investigation aimed at elucidating the role of PQS-induced OMV production during the five stages of biofilm development in *P. aeruginosa.* Here, we report that PQS and OMVs are maximally produced during biofilm dispersion. We further demonstrate that PQS biosynthetic and receptor mutants are deficient in dispersion compared to the wild type. The identified dispersion deficiency was rescued with exogenous PQS, supporting the notion that PQS and PQS-induced OMVs are major contributing factors to *P. aeruginosa* biofilm dispersion. We also demonstrate that purified OMVs possess protease, lipase, and nuclease activities. These results indicate that OMVs may contribute to biofilm dispersion by trafficking enzymes capable of breaking down major EPS components. Through this work, we shed light on a novel role of outer membrane vesicles: the enhancement of biofilm dispersion.

## Results

### PQS production is elevated during dispersion

Although OMVs are ubiquitous in *P. aeruginosa* biofilms (44, 55), their roles and importance in the development of a biofilm remain to be elucidated. PQS is known to promote OMV biogenesis through a biophysical mechanism (30–33), and its synthesis and export are strong indicators of OMV production potential in *P. aeruginosa* (34). The production of PQS is tightly regulated by quorum signaling systems (17, 21, 62, 63), and environmental conditions, such as oxygen availability (61). Due to the heterogeneous nature of biofilm development (64, 65), we hypothesized that PQS-induced OMV production would vary during biofilm progression as nutrient and substrate availability change. Using a continuous flow model, we set out to quantify total PQS production during each stage of biofilm development. Growth stages were determined via microscopic imaging of flow cells using parameters determined by Sauer and coworkers (7). In our system, reversible attachment, irreversible attachment, maturation I, and maturation II were established to occur at 8 h, 24 h, day 3, and day 5, respectively. Dispersion was induced on day five through exogenous addition of the native dispersion cue *cis*-2-decenoic acid (*cis*-DA) Although a *P. aeruginosa* biofilm will naturally produce *cis*-DA and disperse (66), we administered this molecule exogenously in order to synchronize the dispersion event (66, 67). With this study, we found that the highest level of PQS per cell was produced during dispersion (Fig. 1). Concentrations of PQS were normalized to total CFUs and were measured to be 2.7 × 10^−4^ ± 7.2 × 10^−5^, 5.5 × 10^−5^± 2 × 10^−4^, 3.8 × 10^−4^± 1.7 × 10^−4^, 3.0 × 10^−4^ ± 1.5 × 10^−4^ and 3.0 × 10^−3^ ± 4.7 × 10^−4^ µMol per billion CFUs at reversible attachment, irreversible attachment, maturation I, maturation II, and dispersion, respectively (Fig. 1). Statistically significant differences were identified between reversible attachment and dispersion, irreversible attachment and dispersion, maturation I and dispersion, and maturation II and dispersion (one-way ANOVA, Tukey’s post *hoc-test, p* = 0.00020, 0.0015, 0.00080, 0.00020, respectively). In short, a significant increase in PQS was observed in dispersion compared to all other biofilm stages.

**Figure 1.**
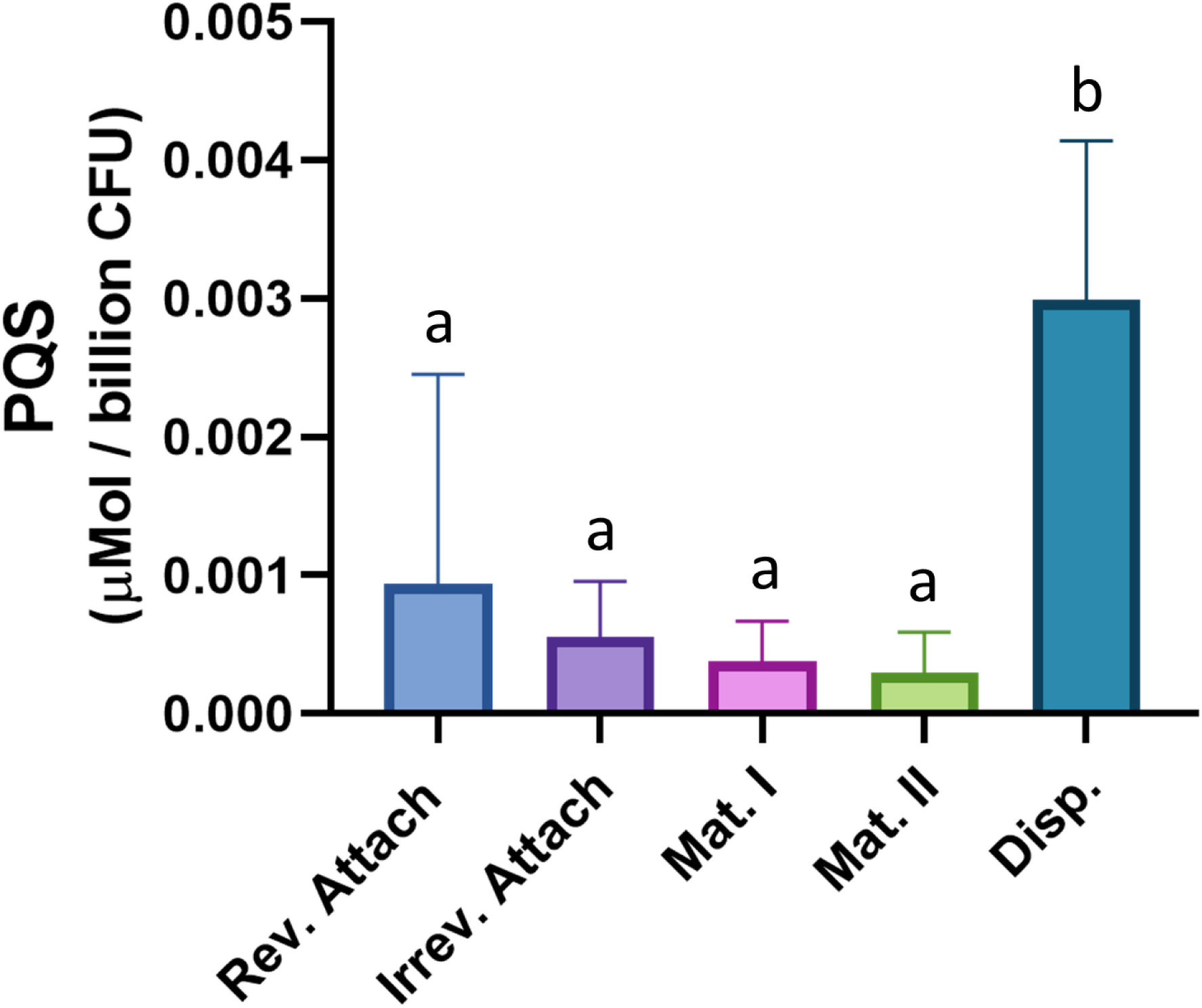
PQS production is elevated during dispersion. PQS was extracted from biofilm tube reactors grown to each of the five stages of development. Measured PQS production was normalized to µMol per billion CFUs. Error bars represent the standard deviation calculated from at least three biological replicates. Statistical significance was assessed by one-way ANOVA followed by Tukey’s *post-hoc* test. Letters above the bars represent significance. Differences between bars that do not share a letter are statistically significant (*p* < 0.05).

### OMV production varies during biofilm development

Following quantification of PQS, OMVs were isolated from the five different biofilm stages and quantified using two independent techniques: OMV protein quantification and nanoparticle tracking analysis (NTA). Modified Lowry assays showed that the highest protein levels were detected in OMV preparations harvested during reversible attachment, irreversible attachment, and dispersion (Fig. 2A). Protein concentrations in OMV pellets were normalized per billion CFUs. The measured values were 94 ± 44, 105 ± 8.5, 11 ± 3.2, 6.5 ± 3.4, and 55 ± 17 µg / billion CFUs during reversible attachment, irreversible attachment, maturation I, maturation II, and dispersion, respectively (Fig. 2A). Statistically significant differences were observed between reversible attachment and maturation I, reversible attachment and maturation II, irreversible attachment and maturation I, irreversible attachment and maturation II, irreversible attachment and dispersion, maturation I and dispersion, and maturation II and dispersion (One-way ANOVA, Tukey’s post *hoc-test, p* = <0.00010, <0.00010, <0.00010, <0.00010, 0.049, 0.025, 0.036, respectively). Quantification via nanoparticle tracking analysis demonstrated that OMV production per cell remained low until the dispersion stage. The particles measured during reversible attachment, irreversible attachment, maturation I, maturation II, and dispersion were 0.44 ± 0.24, 0.69 ± 0.23, 0.74± 0.28, 0.32 ± 0.28, and 2.1 ± 0.37 particles / CFU, respectively (Fig. 2B). Statistically significant differences were identified between reversible attachment and dispersion, irreversible attachment and dispersion, maturation I and dispersion, and maturation II and dispersion (One-way ANOVA, Tukey’s post *hoc-test, p* = <0.00010, <0.00010, 0.00010, <0.00010, respectively). Both quantification techniques showed significantly larger numbers of OMVs present during the dispersion stage compared to the maturation stages. The high level of OMV production during dispersion paralleled enhanced PQS synthesis during this stage. Interestingly, an increase in OMV production during attachment was observed via protein quantification but not through NTA.

**Figure 2.**
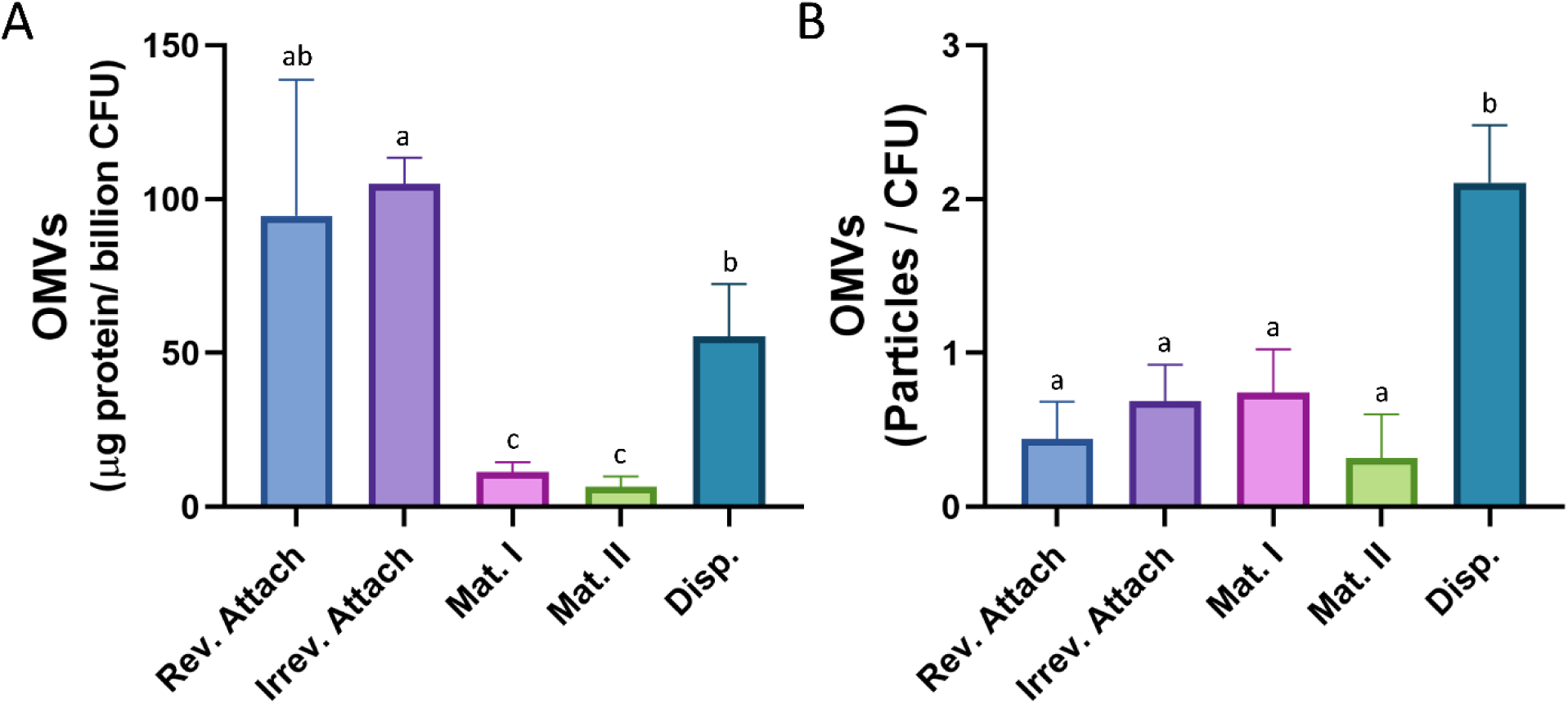
OMV production varies across biofilm developmental stages. OMVs were harvested from each stage of biofilm development and quantified using two different methods. (A) Purified OMVs were quantified by the modified Lowry assay and normalized to µg protein per billion CFUs. (B) Purified OMVs were also quantified using nanoparticle tracking and normalized to CFU. Error bars represent the standard deviation calculated from at least three biological replicates. Statistical significance was assessed by one-way ANOVA followed by Tukey’s *post-hoc* test. Letters above the bars represent significance. Differences between bars that do not share a letter are statistically significant (*p* < 0.05).

### PQS mutants are not deficient in reversible or irreversible attachment

To determine if PQS and/or PQS-controlled phenotypes are involved in the initial stages of *P. aeruginosa* biofilm development, we assessed reversible and irreversible attachment abilities of wild type PA14, Δ*pqsA*, Δ*pqsH*, Δ*pqsE*, and Δ*pqsR*. Crystal violet attachment assays (see methods) were performed at 2 h, 8 h, and 24 h, the former two time points were representative of reversible attachment and the latter was representative of irreversible attachment (7). We found that Δ*pqsA* was not deficient in attachment after 2 or 8 h (Fig. 3A) (Student’s two-tailed *t*-test, *p* = 0.41 and 0.91, respectively) suggesting that quinolones are not involved in reversible attachment. Interestingly, we found that Δ*pqsA* displayed increased attachment after 24 hours (Fig. 3A) (Student’s two-tailed *t*-test, *p* = 0.014). These results indicate that under normal conditions, synthesis of at least one quinolone molecule results in reduced irreversible attachment. Next, we wanted to determine if the observed phenotypes were specifically due to the lack of PQS and PQS-mediated functions. Because Δ*pqsA* is unable to make over 55 different quinolones, we quantified attachment of Δ*pqsH*, which is deficient in synthesis of PQS only (20, 61). We observed no difference in attachment after 2 h or 24 h (Fig. 3B) (One-way ANOVA, *p* = 0.73 and 0.48, respectively). Next, we assessed attachment ability of Δ*pqsE* and Δ*pqsR*, which are unable to induce Rhl-dependent virulence factors (68, 69) and respond to PQS (19), respectively. Reversible (Fig. 3B) and irreversible (Fig. 3C) attachment were unaffected in both mutants (One-way ANOVA, *p* = 0.73 and 0.48, respectively). These results indicate that PQS and PQS-mediated phenotypes do not contribute to the attachment of *P. aeruginosa* to an abiotic surface.

**Figure 3.**
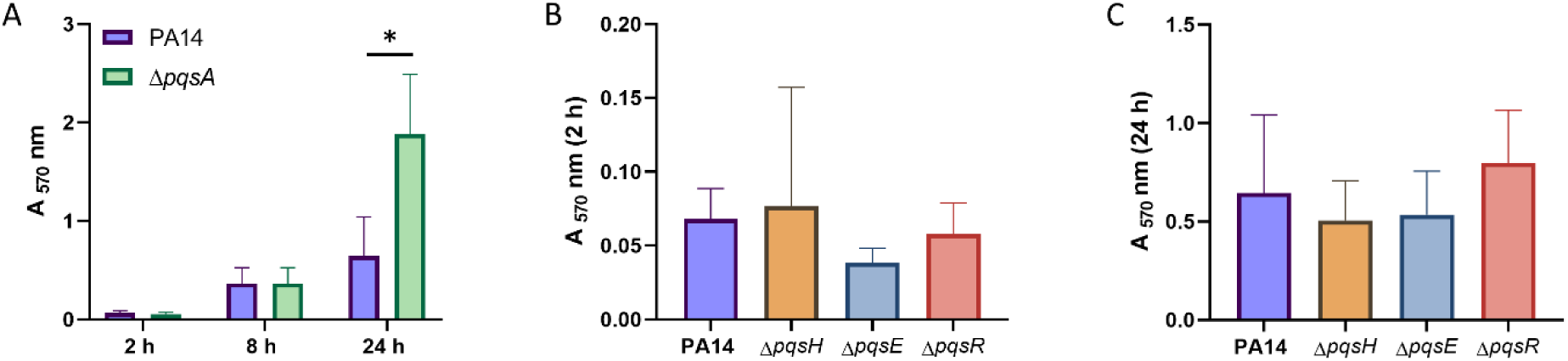
PQS mutants are not deficient in reversible or irreversible attachment. Cultures were grown in 96-well plates, planktonic cells were removed, and attached biomass was quantified using crystal violet staining. (A) PA14 and Δ*pqsA* were grown for 2, 8, and 24 h. (B and C) PA14, Δ*pqsH*, Δ*pqsE*, and Δ*pqsR* were grown for 2 h (B) and 24 h (C). Error bars represent the standard deviation calculated from a minimum of three biological replicates. Statistical significance was determined using Student’s two-tailed *t*-tests for figure 3A and one-way ANOVA for figures 3B and 3C. *, *p* < 0.05.

### ΔpqsA displays diminished biofilm dispersion

Our initial analysis of PQS and OMV production during biofilm development identified that both PQS and OMVs are highly produced during dispersion. To determine if PQS-mediated functions are involved in this stage of development, we quantified dispersion in semi-batch biofilms grown in 24-well plates. On days 4, 5, 6 and 7 after inoculation, microcolonies were observed using light microscopy, and the fraction of microcolonies that had formed central voids, a phenotypic hallmark of the dispersion process in *P. aeruginosa* (7, 9, 67), was determined for PA14 wild type biofilms and for PA14 Δ*pqsA* biofilms. On day 4, little to no dispersion occurred in either strain (Fig. 4A) (Student’s two tailed *t*-test, *p* = 0.87). On days 5, 6 and 7, however, we noted significant differences in microcolony dispersion between the wild type and Δ*pqsA* biofilms (Student’s two-tailed *t*-test, *p* = 0.019, 0.0018, 0.0018, respectively) (Fig. 4A). For subsequent analyses, biofilms were grown until day 6 and analyzed for dispersion. Expression of *pqsA in trans* was able to restore the diminished dispersion phenotype to wild type levels (One-way ANOVA, Tukey’s *post-hoc* test, *p* = 0.63) (Fig. 4B-E).

**Figure 4.**
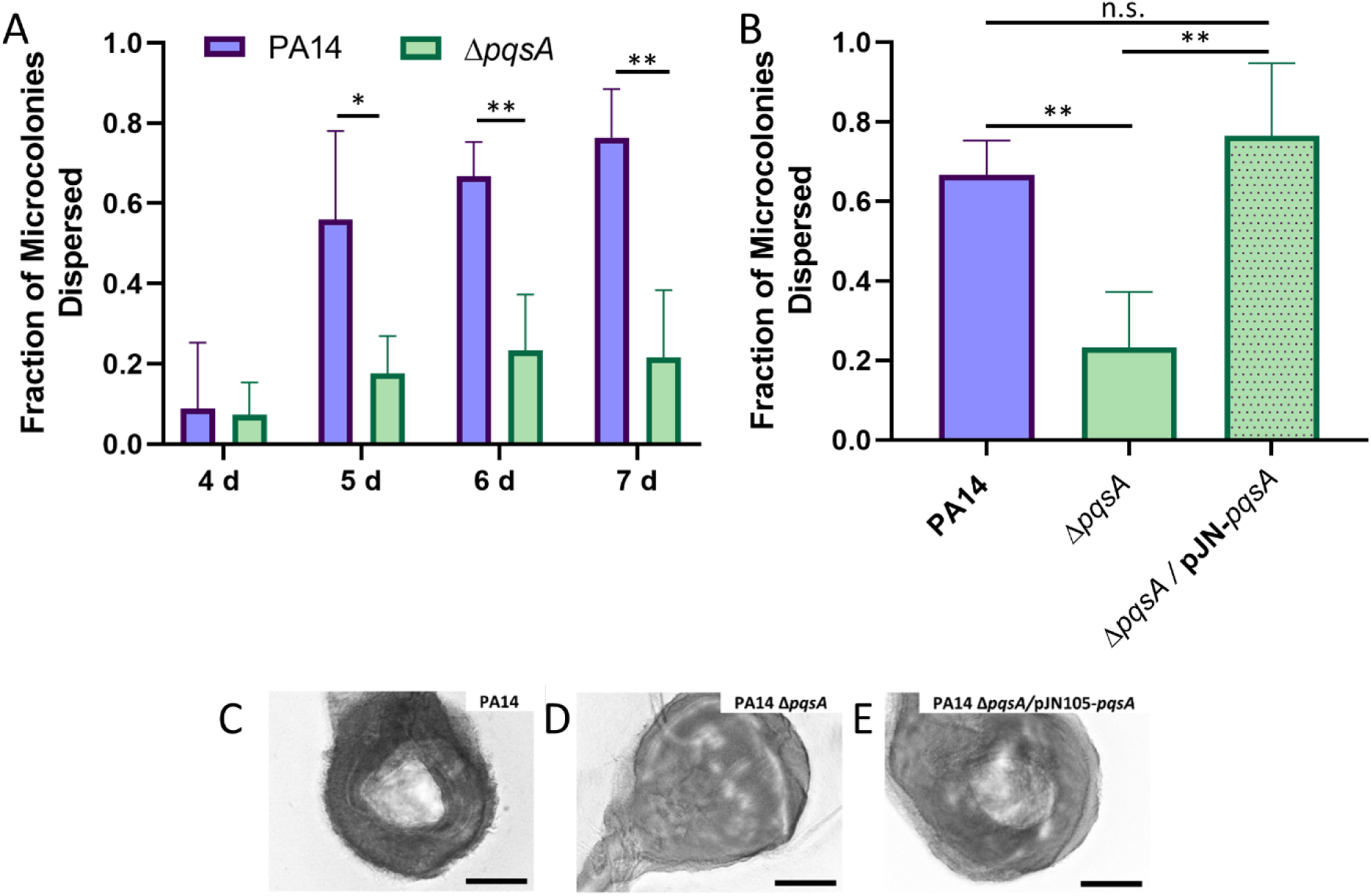
*P. aeruginosa* dispersion is dependent on quinolone biosynthesis. Biofilms were grown in semi-batch cultures in 24-well plates, and the fraction of microcolonies that had dispersed was determined. (A) PA14 wild type and *pqsA* mutant biofilms were assessed for dispersion after 4, 5, 6, and 7 days of growth. (B) Dispersion of the *pqsA* mutant overexpressing the *pqsA* gene was assessed after 6 days of growth and compared to the wild type and *pqsA* mutant. (C-E) Representative images show microcolonies in PA14 wild type (C), PA14 Δ*pqsA* (D), and PA14 Δ*pqsA*/pJN105-*pqsA* (E) biofilms after 6 days of growth. Error bars represent the standard deviation calculated from at least three biological replicates. Scale bars are 100 μm. Statistical significance was determined using Student’s *two-tailed t*-test for figure 4A and one-way ANOVA followed by Tukey’s *post-hoc* test for figure 4B. n.s., *p* > 0.5; *, *p* < 0.05; **, *p* < 0.01.

### P. aeruginosa dispersion is dependent on PQS biosynthesis, but not PqsE

The *pqsA* mutant is deficient in the production of over 55 quinolone molecules (20). For this reason, we were not yet able to conclude whether the inhibition of dispersion was due to a lack of PQS, or a lack of one of the other quinolone molecules. To address this ambiguity, we investigated native dispersion in a *pqsH* mutant. Our results showed that Δ*pqsH* was deficient in dispersion compared to wild type (Fig. 5A). The percentage of microcolonies containing voids in wild type biofilms was 74.68 ± 6.15%, compared to 11.91 ± 3.08% in Δ*pqsH*, suggesting that PQS is specifically responsible for this phenotype (One-way ANOVA, Dunnett’s *post-hoc* test, *p* = 0.0003) (Fig. 5A). However, as PQS is independently involved in both signaling (17) and OMV formation (30, 33, 34), it is unknown whether one or both of these processes are responsible for native levels of dispersion. We also investigated dispersion of a *pqsE* mutant, which produces wild type levels of PQS (21) and OMVs (data not shown), but is deficient in the production of many quorum sensing dependent virulence factors (20). We found that the percentage of microcolonies containing voids in biofilms formed by Δ*pqsE* was 68.69 ± 6.10%, indicating that it disperses at wild type levels (One-way ANOVA, Dunnett’s *post-hoc* test, *p* = 0.86) (Fig. 5A). This suggests that a non-signaling-dependent function of the PQS system, such as OMV production, is likely responsible for the diminished dispersion phenotype in the Δ*pqsA* and Δ*pqsH* mutants. We also investigated dispersion in the *pqsR* mutant, which displays reduced production of both PQS and OMVs (21, 30). The percentage of microcolonies containing voids in biofilms formed by Δ*pqsR* was 37.48 ± 18.97% and significantly lower than wild type (One-way ANOVA, Dunnett’s *post-hoc* test, *p* = 0.0065) (Fig. 5A). The reduced dispersion phenotype of the Δ*pqsH* and the Δ*pqsR* mutants was restored to wild type levels through genetic complementation (Fig. 5B). The percentages of microcolonies containing voids in biofilms formed by PA14 / pJN105, Δ*pqsH* / pJN105-*pqsH*, and Δ*pqsR* / pJN105-*pqsR* strains were 73.24 ± 12.35%, 85.20 ± 4.92%, and 81.80 ± 9.92%, respectively (Fig. 5B). These data suggest that PQS-induced OMV production plays a significant role in *P. aeruginosa* biofilm dispersion.

**Figure 5.**
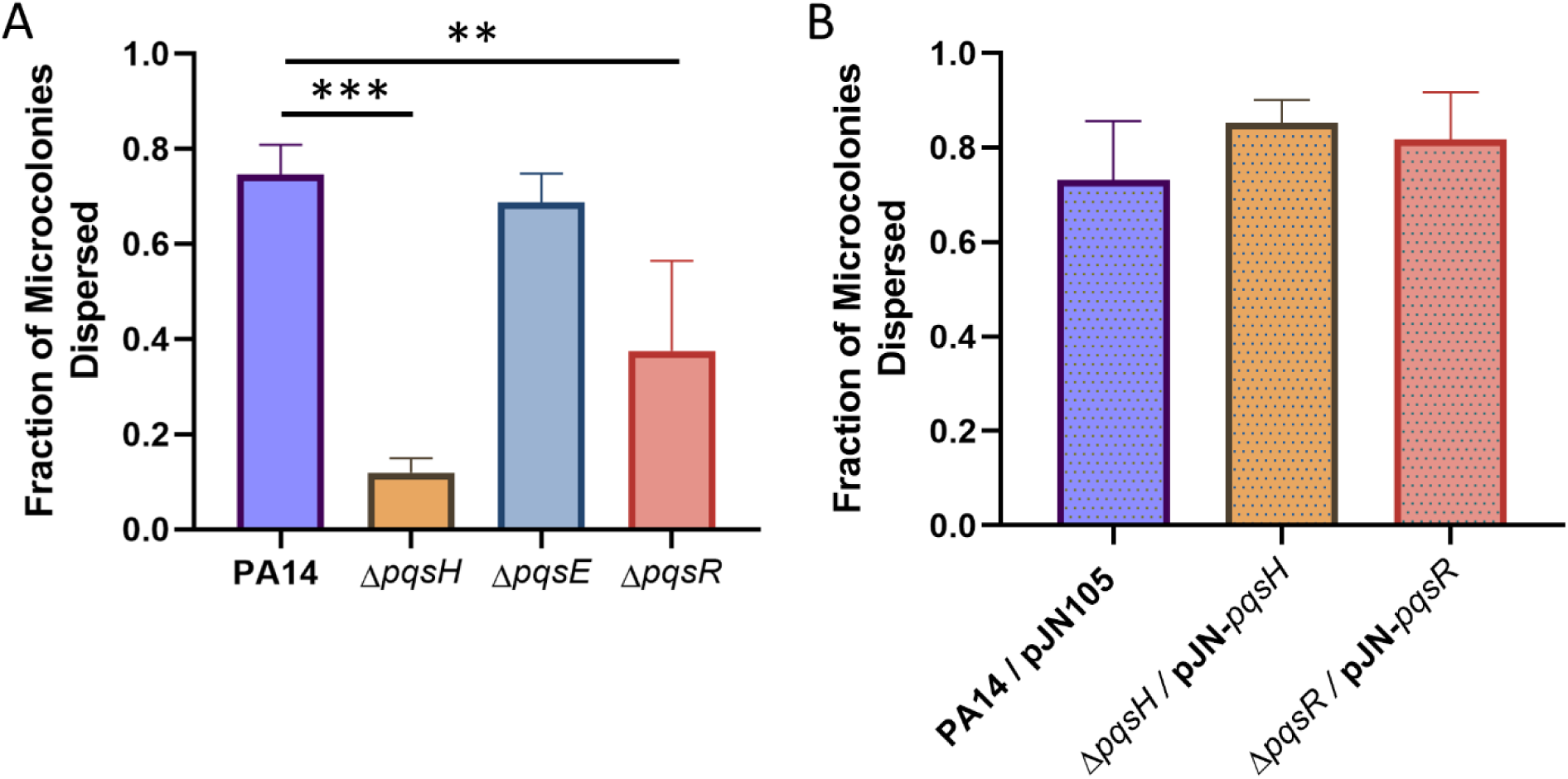
Production of PQS specifically restores native biofilm dispersion. Biofilms were grown in semi-batch cultures in 24-well plates for 6 days. (A) The fraction of microcolonies dispersed was found for PA14 wild type biofilms as well as Δ*pqsH*, Δ*pqsE*, and Δ*pqsR* biofilms. (B) Overexpression of the missing genes in the mutant backgrounds restored the dispersion phenotype that was diminished in Δ*pqsH* and Δ*pqsR* biofilms. Bars represent the standard deviation calculated from at least three biological replicates. Statistical significance was analyzed by one-way ANOVA followed by Dunnett’s *post-hoc* test. **, *p* < 0.01; ***, *p* < 0.001.

### Exogenous PQS restores dispersion in the ΔpqsR mutant

To confirm whether PQS modulates dispersion through an OMV-dependent mechanism, exogenous PQS was administered to a Δ*pqsR* biofilm and dispersion efficiency was quantified. PQS-induced OMV production has been shown to be driven by a biophysical mechanism that is not signaling dependent (31–33). The exogenous addition of PQS to a Δ*pqsR* biofilm restored dispersion to wild type levels (One-way ANOVA, Tukey’s *post-hoc* test, *p* = 0.72) (Fig. 6). Microcolony void formation increased from 60.65 ± 3.12% to 77.09 ± 6.94% (One-way ANOVA, Tukey’s *post-hoc* test, *p* = 0.024) (Fig. 6). This indicates that PQS modulates dispersion using an OMV-dependent mechanism that is separate from the PQS signaling network.

**Figure 6.**
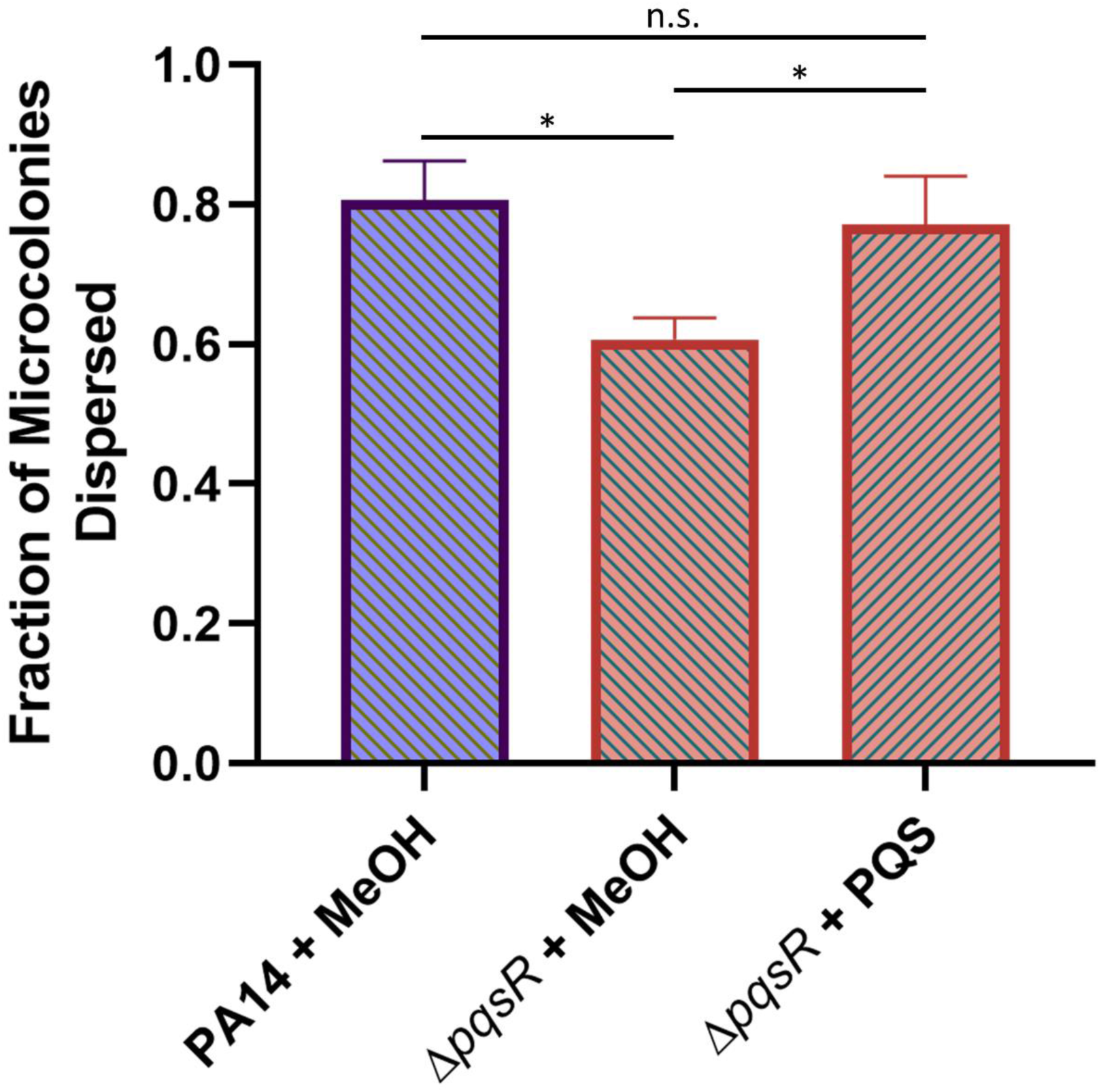
Exogenous PQS rescues Δ*pqsR* dispersion defect. PA14 wild type and Δ*pqsR* biofilms were grown in semi-batch cultures in 24-well plates for 4 days. For the following 2 days, the medium was exchanged every 12 hours with fresh medium containing 40 μM PQS (+ PQS), or an equivalent amount of methanol (+ MeOH, vehicle control). Dispersion efficiency was then quantified for the strains under each condition. Error bars represent the standard deviation calculated from at least three biological replicates. Statistical significance was analyzed by ANOVA followed by Tukey’s *post-hoc* test. n.s., *p* > 0.5; *, *p* < 0.05.

### OMVs contain enzymes capable of degrading the biofilm matrix

Together, our results indicate that PQS-induced OMVs contribute to the dispersion of *P. aeruginosa* biofilms; however, the exact role the vesicles play during this developmental stage is unknown. Various studies have demonstrated that degradation of extracellular polymeric substances (EPS) of the biofilm matrix, such as polysaccharides, proteins, glycolipids, and eDNA, is a requirement for dispersion (reviewed in (15)). Degradative enzyme activity towards these matrix components has been shown to induce dispersion in both Gram-positive and Gram-negative organisms (15, 70–76) Previous OMV proteomic analyses have identified several proteins packaged within vesicles that were predicted to have degradative activity (77, 78). Therefore, we hypothesized that OMVs may contribute to dispersion through EPS degradation. To test this hypothesis, we assessed whether purified *P. aeruginosa* OMVs were capable of degrading skim milk, tributyrin, and DNA to assess protease, lipase, and DNase activity, respectively. In order to acquire sufficient material for these analyses, planktonic OMVs were used. Addition of OMVs to skim milk agar resulted in the formation of a 119.8 ± 36.1 mm^3^ zone of clearing, while the addition of vehicle control (MV buffer only) to skim milk agar resulted in the formation of a 0.1 ± 8.6 mm^3^ zone of clearing (Student’s two-tailed *t*-test, *p* = 0.0007) (Fig. 7A). This suggests that OMVs contain enzymes that have protease activity. The addition of OMVs to tributyrin agar resulted in the formation of a 211.1 ± 24.1 mm^3^ zone of clearing *versus* the vehicle control that produced a 25.9 ± 11.2 mm^3^ zone of clearing (Student’s two-tailed *t*-test, *p* < 0.0001) (Fig. 7B). This suggests that OMVs also contain enzymes that have lipase activity. Finally, the addition of OMVs and vehicle control to DNase agar resulted in the formation of 182.1 ±85.5 mm^3^ and 21.3 ±16.3 mm^3^ zones of clearing, respectively (Student’s two-*tailed t-test, p* = 0.010) (Fig. 7C). This indicates that OMVs carry enzymes with DNase activity. Overall, these data support the idea that OMVs contribute to biofilm dispersion by packaging and delivering enzymes with EPS degrading abilities.

**Figure 7.**
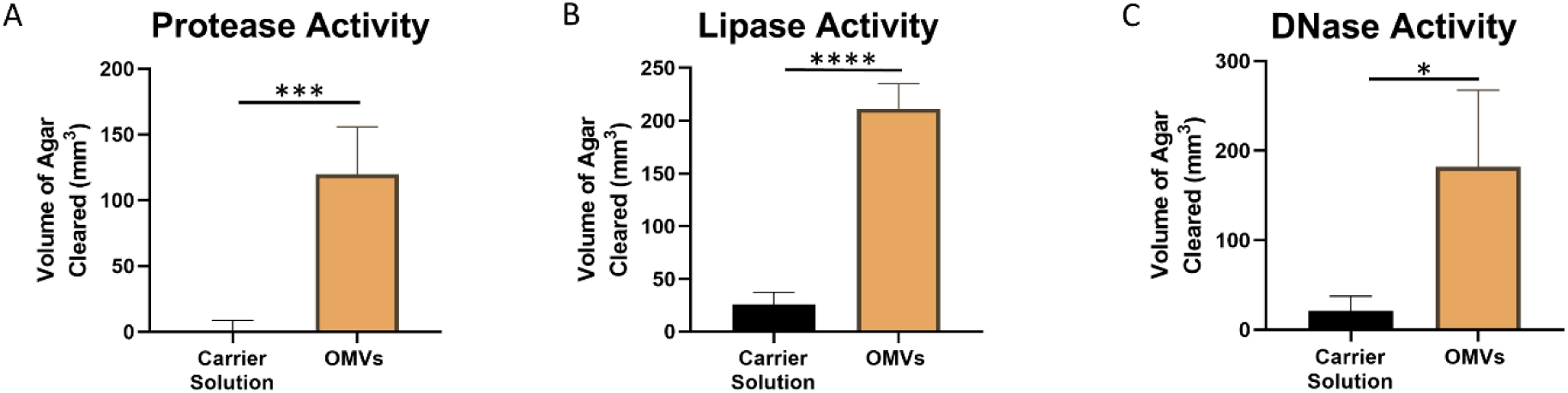
Purified OMVs display EPS-degrading activities. OMVs were harvested, washed with, and resuspended in MV buffer, and added to wells punched into different types of agar. (A) Skim milk agar was used to assess protease activity. (B) Tributyrin agar was used to assess lipase activity. (C) DNase agar to assess DNase activity. Error bars represent the standard deviation calculated from three biological replicates. Significance was assessed using Student’s two-tailed *t*-tests. *, *p* < 0.05; ***, *p* < 0.001; ****, *p* < 0.0001.

## Discussion

Biofilms have become a major health and economic concern due to their prevalence and recalcitrance. *P. aeruginosa* is a leading cause of nosocomial infections (79), as well as increased morbidity and mortality in cystic fibrosis patients (80). Virulence and pathogenesis in this organism are largely regulated by quorum sensing signals (81). PQS is one such signal that controls the production of virulence factors (17) but is also known to induce production of OMVs (30, 33, 61). OMVs represent a dedicated trafficking system that delivers virulence factors (47, 82), while also carrying cargo able to degrade antibiotics (45), lyse neighboring bacteria (30, 47, 55), and enable cell-to-cell communication (30). Several groups have demonstrated that OMV production is prevalent in biofilms (44, 54, 55). However, the biogenesis and function of OMVs during biofilm development remains poorly understood, as most of what is known about OMVs comes from studies of planktonic bacteria. The present study set out to elucidate the role of PQS-induced OMV production in *P. aeruginosa* during the five distinct stages of biofilm development.

PQS is an excellent predictor of OMV production (30, 34) and studies have consistently shown that a block in PQS synthesis (whether genetic or environmental) results in dramatically reduced OMV formation (30, 55, 61). Although extracellular vesicles have been observed in the absence of PQS (54, 55), their origins and composition are uncertain, and they are frequently mixed-composition vesicles resulting from cellular disintegration. For this reason, we were surprised to measure high levels of OMVs during reversible and irreversible attachment using protein-based quantification, despite low PQS concentrations (Fig. 1 and Fig. 2). High levels of OMV production during these initial stages measured by Lowry assay were not corroborated by nanoparticle tracking analysis, suggesting that the protein detected in these OMV preparations was not representative of OMV concentration but likely the result of non-OMV-related protein components. As a result, we predicted that PQS and OMVs were not significant effectors of reversible and irreversible attachment. This notion was supported by our crystal violet attachment assays, which demonstrated that Δ*pqsA*, Δ*pqsH*, Δ*pqsR*, and Δ*pqsE* mutants had wild type levels of reversible attachment (Fig. 3). It is notable, however, that several studies have identified an increase in biofilm formation when OMV production is stimulated (22, 51, 83, 84). Kang *et al*. (23) described that *pqsA*, but not *pqsH* or *pqsE*, was required for early biofilm attachment under static conditions. Others have reported that PQS, and possibly OMVs, were more important in later maturation stages (26, 27, 85). In contrast, Ionescu *et. al.* showed in *Xylella fastidiosa* that OMV production *inhibited* bacterial attachment to plant surfaces, increased bacterial motility, and enhanced plant mortality (86). In the face of these conflicting reports, it is interesting that we found the *pqsA* mutant had increased irreversible attachment *versus* wild type at 24 hours (Fig. 3A). During early biofilm development attachment is required. Therefore, it might be beneficial for *P. aeruginosa* to reduce PQS production at this time to avoid potential interference of PQS-induced OMVs with cell attachment. Regardless, it is evident that the role of OMVs in early-stage biofilm development remains unclear and will require further studies to elucidate.

During maturation I and II, we saw that both PQS and OMV production were relatively low (Fig. 1 and Fig. 2). Allesen-Holm *et al.* described PQS’ role in the development of three-dimensional microcolony architecture (26). They proposed that PQS induced prophage-mediated cell lysis, resulting in eDNA release and increased biofilm formation (26). A separate study by Tettman *et al.* showed that enzymatic degradation of PQS resulted in increased iron availability and enhanced biofilm formation for early and mature biofilms (87). The latter report aligns with our observations and offers an explanation as to why cells might reduce PQS production during biofilm maturation. It is important to note that although PQS production was reduced during maturation in our study, it was not eliminated. The same was true for OMV production. It is likely that baseline levels of PQS are important for PQS-mediated cell lysis and eDNA release while reduced numbers of OMVs may carry out structural or transportation roles. At this developmental stage, elevated levels of PQS and PQS-induced OMVs could even have negative effects on biofilm development, as OMVs have been predicted to contain degradative enzymes (77, 78), which could break down major components of the EPS.

While our results suggest that PQS and OMVs may play only minor (or undetermined) roles during attachment and maturation, they highlight a major increase in production of both factors upon the initiation of biofilm dispersion (Fig. 1 and Fig. 2). This observation led us to speculate that PQS and PQS-induced OMVs are important for proper dispersion of *P. aeruginosa* biofilms. To test this hypothesis, we analyzed microcolony dispersion frequencies for four mutants: Δ*pqsA*, Δ*pqsH*, Δ*pqsR*, and Δ*pqsE.* Biosynthetic (*pqsA, pqsH*) and receptor (*pqsR*) mutants dispersed at much lower frequencies than wild type (Fig. 4 and Fig. 5). Because the Δ*pqsA* and Δ*pqsH* mutants were similarly impaired in dispersion, we can conclude that PQS, specifically, is required (i.e. not any of the other alkyl-quinolones lost in the Δ*pqsA* mutant). Rescue of the Δ*pqsR* phenotype by exogenous PQS demonstrated that the *physical presence* of PQS was required, rather than signaling through its receptor (Fig. 6). The importance of a non-signaling function of PQS is further supported by the fact that the *pqsE* mutant showed no deficiency in dispersion, confirming that signaling downstream of PqsR is also not involved in this phenotype (Fig. 5). Together, these results demonstrate that PQS modulates *P. aeruginosa* dispersion in a signaling-independent manner. Our final experiments led us to propose that PQS-induced OMVs, which are formed through a signaling-independent biophysical mechanism (30, 33), promote dispersion by carrying EPS degrading enzymes.

EPS degradation is a fundamental requirement for dispersion (15). Enzymes with matrix degradative activity have been described to induce dispersion in mature biofilms in several organisms (15, 71–76, 88). The effectiveness of DNaseI at dispersing biofilms has even led to its adoption as a treatment for biofilm infections in the lungs of cystic fibrosis patients (89). Previous studies have shown that *P. aeruginosa* OMVs have autolysin (47, 48), and protease (44, 90) activity, and that these OMVs can associate with and lyse bacterial sacculi (47). These findings support the proposition that OMVs carry degradative enzymes. Here, we report that purified OMVs possess protease, lipase, and DNase activity (Fig. 7). A recent study by Esoda and Kuehn also found that OMVs traffic the *P. aeruginosa* peptidase, PaAP, and can deliver the peptidase to 1-hour old *P. aeruginosa* and *K. pneumoniae* biofilms grown on A549 tissue culture cells, resulting in decreased biofilm biomass (91). Others have provided evidence that proteases are required for dispersion in *S. aureus* biofilms (71) and *P. putida* biofilms (73). In *P. aeruginosa*, eDNA degradation has been shown to result in biofilm disaggregation (26, 92) and recent work by Cherny and Sauer showed that eDNA degradation is required for dispersion of *P. aeruginosa* (72). In *P. acnes*, secreted lipases have also been demonstrated to enhance the dispersion response (93). Delivery of these degradative enzymes using OMVs may increase the enzymes’ efficacy, facilitate specific targeting to sites of degradation, and reduce potential deactivation of the enzymes while in transit. Bomberger *et al.* demonstrated that the CFTR inhibitory factor (Cif) produced by *P. aeruginosa* was orders-of-magnitude more potent when delivered within OMVs (82). We therefore propose that PQS-induced OMVs enhance biofilm dispersion by delivering and potentially enhancing the activity of enzymes required for EPS degradation.

Previous studies have identified the importance of PQS in biofilm formation (26, 27) and demonstrated the presence of OMVs within biofilms (44). However, a comprehensive study that analyzed the effect of these two factors at each stage in biofilm development had not been conducted prior to this work. Here, we report that PQS and OMVs are not produced consistently during biofilm development; specifically, we identified low (or variable) concentrations of PQS and OMVs during attachment and maturation stages but high concentrations during dispersion. Additionally, we showed that attachment is likely not affected by the absence of PQS and PQS-mediated factors, whereas the absence of PQS significantly reduces dispersion of *P. aeruginosa* biofilms. Finally, we demonstrated that OMVs have the capability to breakdown extracellular DNA, lipids, and proteins – all major components of the biofilm EPS matrix. With this work we identified PQS and PQS-induced OMVs as novel regulators of biofilm dispersion. Because dispersed cells are significantly more susceptible to antimicrobials (94–96), it has been considered that dispersion agents in combination with antimicrobials could provide a potent antibiofilm therapy. Therefore, PQS and PQS-induced OMVs may provide novel avenues to create better treatment strategies against recalcitrant biofilm infections.

## Materials and Methods

### Strains, growth conditions, and media

All experiments were carried out using *P. aeruginosa* strains described in Table 1. The Δ*pqsE* and Δ*pqsR* clean-deletion mutant strains were constructed using the pEX18gm suicide vector (97), and *pqsE* and *pqsR* were overexpressed in their respective mutant backgrounds using the pJN105 vector (98). Primer sequences used for construction of the vectors can be found in Table S1. Biofilm tube reactors were inoculated as described below. Planktonic cultures were inoculated to an OD_600_ of 0.01 and grown at 37 °C with shaking at 250 rpm. Planktonic cultures were grown in Lysogeny Broth (LB) or brain heart infusion medium (BHI). Planktonic cultures of strains carrying the pJN105 vector were grown in the presence of gentamicin (50 µg/mL), while biofilm cultures of the same strains were not.

**Table 1.** Bacterial strains and plasmids used in this study.

| Strain or Plasmid | Description | Source or Reference |
| --- | --- | --- |
| **Strains:** |  |  |
| *E. coli* |  |  |
| DH5α | F– Φ80lacZΔM15 Δ(lacZYA-argF) U169 recA1 endA1 hsdR17 (rK–, mK+) phoA supE44 λ– thi-1 gyrA96 relA1 | (103) |
| *P. aeruginosa* |  |  |
| PA14 | Wild type *P. aeruginosa* strain | (104) |
| Δ*pqsA* | *pqsA* clean deletion in PA14 background | Kind gift of Marvin Whiteley |
| Δ*pqsH* | *pqsH* clean deletion in PA14 background | (105) |
| Δ*pqsE* | *pqsE* clean deletion in PA14 background | This study |
| $\Delta pqsR$ | *pqsR* clean deletion in PA14 background | This study |
| **Plasmids** |  |  |
| pEX18gm | Gm<sup>R</sup>; suicide plasmid for gene replacement in *P. aeruginosa* | (97) |
| pEX18gm-*pqsE* | Gm<sup>R</sup>; pEX18gm-derived vector for clean-deletion of *pqsE* | This study |
| pEX18gm-*pqsR* | Gm<sup>R</sup>; pEX18gm-derived vector for clean-deletion of *pqsR* | This study |
| pJN105 | Gm<sup>R</sup>; *araC*-*pBAD* expression vector | (98) |
| pJN105-*pqsA* | Gm<sup>R</sup>; pJN105-derived *pqsA* overexpression vector | (55) |
| pJN105-*pqsH* | Gm<sup>R</sup>; pJN105-derived *pqsH* overexpression vector | This study |
| pJN105-*pqsR* | Gm<sup>R</sup>; pJN105-derived *pqsR* overexpression vector | This study |
| pCR 2.1 | Amp<sup>R</sup>; Kan<sup>R</sup>; TA-cloning vector | Invitrogen |
| pRK2013 | Km<sup>R</sup>; Helper plasmid used for triparental mating | (106) |

### Biofilm growth

Biofilms were grown in both continuous and semi-batch culture systems. For continuous culture, biofilms were grown in size 14 Masterflex silicone tubing (Cole Parmer) as previously described (7, 99). Cultures were inoculated under static conditions and allowed to attach for 1 h prior to initiation of flow. Biofilms were grown at 22°C in 5% LB medium under a constant flow rate of 0.18 mL/min until desired stage of biofilm growth; 8 h for reversible attachment, 24 h for irreversible attachment, 3 days for maturation I, 5 days for maturation II (as determined previously (7) and in this study by microscopic flow cell images). To validate developmental stages, biofilms were grown under identical conditions in BioSurface Technologies flow cells and visualized by brightfield microscopy. Biofilms were harvested from continuous culture systems using the rolling pin method (7). Mature biofilms were collected into sterile saline (1mL / line). For stage 5, biofilm dispersion, 5% LB with or without the native dispersion induction molecule *cis*-2-decenoic acid (310 nM) was administered to five-day old biofilms. Biofilms were incubated with either treated or untreated medium under static flow for 1 hour (66, 67). Following induction, dispersed cells in the bulk liquid were collected under native flow, leaving attached biofilm cells behind in the tubing. To quantify if a dispersion event occurred, OD_600_ measurements were taken of the collected bulk liquid from the treated sample and compared to the untreated sample.

Semi-batch biofilms for dispersion analyses were cultured in 24-well plates as previously described (67) with minor modifications. Briefly, wells were inoculated with 500 µL of culture adjusted to an OD_600_ of 0.01 in 20% LB. Plates were incubated at 37°C with shaking at 250 rpm at a 30° angle for 24 h. Media was then replaced with 250 µL of 20% LB medium and returned to the incubator under the same conditions. Media changes were repeated every 12 h for up to 7 days. For chemical complementation experiments, strains were inoculated and grown as described above for the first 4 days. From 4 days post inoculation to 6 days post inoculation, media was changed with 20% LB containing 40 μM PQS or 20% LB containing and equivalent amount of the carrier solution (methanol) every 12 hours.

### PQS extraction and quantification

PQS was extracted from biofilms harvested at each stage of development. Biofilms were homogenized to reduce aggregation and PQS was extracted using 1:1 acidified ethyl acetate as previously described (34, 55, 61, 100). The organic phase was separated and dried under nitrogen. Samples were resuspended in optima grade methanol and spotted onto straight-phase phosphate-impregnated TLC plates that had been activated at 100°C for 1 h. PQS was visualized by intrinsic fluorescence after excitation under long-wave UV light. Digital images were captured and analyzed using a BioRad ChemiDoc XRS system and Image Lab densitometry software. PQS concentration values were normalized to total CFUs.

### OMV isolation and quantification

OMVs were isolated from harvested biofilms as previously described (55). Biofilms were homogenized to reduce aggregation and preparations were centrifuged at 16,000xg for 10 min at 4°C to remove cells. The supernatant was then passed through a 0.45 µm polyethersulfone filter to remove any remaining cells. OMVs were pelleted and purified from the supernatant using a Thermo Scientific S50-A rotor (50,000 rpm for 1.5 h) and resuspended in 500 µL of sterile MV buffer (50 mM Tris, 5 mM NaCl, 1 mM MgSO4, pH 7.4) (34, 55).

OMVs were then quantified by both modified Lowry protein assay (Thermo) (101) and nanoparticle tracking analysis (NTA) (34, 55, 102). The modified Lowry assay was performed following manufacturer’s instructions. Purified vesicles were diluted to obtain 20-100 particles per frame and analyzed using a NanoSight NS3000 system (camera level 12 and gain of 1) and corresponding software (NTA 3.1). Total protein and OMV particle values were normalized to total CFUs in the original sample.

### Crystal violet attachment assays

To assess attachment, 96-well plates were inoculated with 200 μL of culture in LB at an OD_600_ of 0.01. The plates were then incubated at 37°C shaking at 250 rpm for 2, 8, or 24 h. Biomass was quantified by crystal violet (CV) staining. Supernatant was removed from wells and replaced by 200 µL DI water. 50 μL of 0.1% CV in DI water was then added to each well, and plates were incubated for 15 minutes at 37°C with shaking at 250 rpm. Following staining, wells were washed 4 times with DI water to remove any unattached cells and unbound CV. Plates were then blotted vigorously onto paper towel and allowed to dry. Once dry, 200 μL of 95% ethanol was added to each well and the plate was incubated for 10 minutes at 37°C with shaking at 250 rpm to solubilize the CV. The absorbance of each well was then read at 570 nm.

### Assessment of dispersion phenotype in 24-well microtiter plates

Biofilms were grown as described above for up to 7 days, and native dispersion was assessed as previously described (9, 67). Briefly, biofilm microcolonies were observed by transmitted light using an Olympus BX60 microscope and a 20 × UPlanF Olympus objective. Images were captured using a ProgRes CF camera (Jenoptik, Jena, Thuringia, Germany) and processed with ProgRes CapturePro 2.7.7 software. Dispersion efficiency was quantified by determining the percentage of microcolonies that had developed an interior void. For each biological replicate, biofilms were grown in 2 to 4 wells of a 24-well plate, and all microcolonies that had formed in these biofilms were analyzed for dispersion. The total number of microcolonies analyzed for each strain and condition are presented in Supplemental table 2.

### Analysis of degradative enzyme presence in OMVs

In order to acquire enough material for enzymatic analysis, OMVs were harvested from planktonic cultures as described above. OMV preparations were quantified using NTA and diluted to 2×10^11^ particles/mL in MV buffer. 180 μL of OMVs were then added to wells punched in agar using a method described previously (93). Agar plates impregnated with protein, lipid or DNA were prepared, and wells were punched within the agar using the wide end of a 1000 μL pipette tip. Each 100 mm diameter petri dish used contained 25 mL of an agar solution. For proteomic analysis, milk agar plates were prepared (2.5 g/L skim milk (BD) and 15 g/L agar (BD)). For these plates, skim milk and agar were autoclaved separately, cooled to 50°C, and then mixed together prior to pouring plates. For lipase analysis, 50% tributyrin agar was used (11.5 g/L Tributyrin HiVeg Agar Base (HiMedia), 5 mL/L Tributyrin (TCI), 7.5 g/L agar (BD)). Specifically, the agar was boiled in water, tributyrin was added, and the mixture was homogenized in a blender for approximately 20 seconds to ensure effective dispersal of the hydrophobic tributyrin throughout the medium. Once autoclaved, this agar was stirred while cooling to approximately 60°C, and the plates were then poured. For DNase analysis, DNase plates were prepared (21 g/L Difco™ DNase test agar with methyl green (BD), 7.5 g/L agar (BD)). After addition of OMVs into the punched wells, plates were sealed with parafilm and incubated at 37°C for 24 h prior to measuring the diameter of the zone of clearing.

### Statistical Analysis

Statistical analyses were performed as described in figure legends and carried out in GraphPad Prism 8.

## Acknowledgements

We thank former First-year and Summer Research Immersion Program students, Maria Carlucci, Ana Conceicao, Wilmer Estevez, Channelle Farquharson, Avery Hoda, Crystal Huang, Nadia Mirza, Laura Oliveira, Sonny Pohar, Sarah Pokrzywa, Kayla Principe, Michael Toledano, Antonio Torlentino, Kyra Yanusas, and former Schertzer Lab students Alexis Gursky, Nicole Radova and Nikki Naim for their contributions to this project. We also thank David Davies and Amanda Zdimal for their assistance with the degradative enzyme assays.

This work was supported in part by grants from the NIH (1R21AI121848 and 1R15GM135862 to J.W.S.), the Research Foundation of SUNY (to J.W.S), to Binghamton University from the Howard Hughes Medical Institute (HHMI) through the Precollege and Undergraduate Science Education Program, and from the New York State Regional Economic Development Council for the First-year and Summer Research Immersion Programs. The content is solely the responsibility of the authors and does not necessarily represent the official views of the National Institutes of Health.

